# Characterization and Structural Prediction of ORF10, ORF7b, ORF7a, ORF6, Membrane Glycoprotein, and Envelope Protein in SARS-CoV-2 Bangladeshi Variant through Bioinformatics Approach

**DOI:** 10.1101/2021.10.01.452232

**Authors:** Pinky Debnath, Umama Khan, Md. Salauddin Khan

**Affiliations:** Chemical Biotechnology Department, Technical University of Munich, Schuigasse 22, D-94315 Straubing, Germany; Biotechnology and Genetic Engineering Discipline, Khulna University, Khulna-9208, Bangladesh; Statistics Discipline, Khulna University, Khulna-9208, Bangladesh

**Keywords:** SARS-CoV-2, ORF proteins, Membrane and Envelope protein, Bangladeshi Covid-19 variant, Structural prediction, Bioinformatics

## Abstract

The acute respiratory disease induced by the severe acute respiratory syndrome-coronavirus-2 (SARS-CoV-2) has become a global epidemic in just less than a year by the first half of 2020. The subsequent efficient human-to-human transmission of this virus eventually affected millions of people worldwide. The most devastating thing is that the infection rate is continuously uprising and resulting in significant mortality especially among the older age population and those with health co-morbidities. This enveloped, positive-sense RNA virus is chiefly responsible for the infection of the upper respiratory system. The virulence of the SARS-CoV-2 is mostly regulated by its proteins like entry to the host cell through fusion mechanism, fusion of infected cells with neighboring uninfected cells to spread the virus, inhibition of host gene expression, cellular differentiation, apoptosis, mitochondrial biogenesis, etc. But very little is known about the protein structures and functionalities. Therefore, the main purpose of this study is to learn more about these proteins through bioinformatics approaches. In this study, ORF10, ORF7b, ORF7a, ORF6, membrane glycoprotein, and envelope protein have been selected from a Bangladeshi Corona-virus strain G039392 and a number of bioinformatics tools (MEGA-X-V10.1.7, PONDR®, ProtScale, ProtParam, SCRIBER, NetSurfP v2.0, IntFOLD, UCSF Chimera, and PyMol) and strategies were implemented for multiple sequence alignment and phylogeny analysis with 9 different variants, predicting hydropathicity, amino acid compositions, protein-binding propensity, protein disorders, 2D and 3D protein modeling. Selected proteins were characterized as highly flexible, structurally and electrostatically extremely stable, ordered, biologically active, hydrophobic, and closely related to the proteins of different variants. This detailed information regarding the characterization and structure of proteins of SARS-CoV-2 Bangladeshi variant was performed for the first time ever to unveil the deep mechanism behind the virulence features and also, this robust appraisal paves the future way for molecular docking, vaccine development targeting these characterized proteins.

## 1. Introduction

In December 2019, the whole world was stunned by the outbreak of unknown cause pneumonia, which was originated from Wuhan, Hubei province of China. And then, by Jan 7, 2020, Chinese scientists have screened a novel Coronavirus (CoV) mainly responsible for the infection of the upper respiratory system from patients in Wuhan [1]. The ensuing proficient human-to-human transmission of the virus ultimately affected millions of people worldwide. Since July 2021, there were 183 million confirmed cases and 3.97 million people have died around the world by this devastating virus. The most devastating thing is that the infection rate is continuously uprising which is resulting in significant mortality especially among the older age population and those with health co-morbidities.

Coronaviruses are very tiny in size (diameter, 65-125 nm) consist of a single-strand RNA as nucleic material [2,3]. Along with RNA, this particular virus consists of 12 different proteins such as nonstructural proteins (ORF1a and ORF1b) at the 5′-end, structural proteins (spike surface glycoprotein (S), envelope (E), matrix (M), and nucleocapsid (N)) and multiple lineage-specific accessory proteins (ORF3a, ORF6, ORF7b, ORF8, and ORF10) at the 3′-end [4]. Although these proteins are basically involved in host receptor recognition, attachment, and entry into host cells, very slight is recognized about these protein structures and specific functionalities. The ORF10 protein is found upstream of the 3’-untranslated region (3’-UTR), apparently encodes for a protein of 38 amino acids long [5]. The ORF7b protein is a presumed viral accessory protein encoded on subgenomic (sg) RNA 7 [6], whereas, ORF7a possessed a distinctive immunoglobulin (Ig)-like domain with a 15-a.a single peptide sequence at its N terminus, an 81-a.a luminal domain, a 21 a.a transmembrane domain, and a short C-terminal tail [7]. Also, the SARS-CoV ORF6 is characteristically between 42 to 63 amino acids in length and by transcribing into mRNA6 encodes SARS 6 protein [8]. Moreover, the membrane glycoprotein is found abundantly and plays the main role in virion assemble, morphogenesis, and, also, define the shape of the viral envelope [9,10]. Lastly, the envelope proteins are short-chain polypeptide with a single α-helical transmembrane domain that can produce homopentametric ion channels (IC) [11].

Though Bangladesh is not exempt from the severe outbreak of the Coronavirus, a large number of Bangladeshi strains also have been identified. On 8 March 2020, in Bangladesh, SARS-CoV-2 was reported for the very first time. A new strain was acknowledged on January 2021, from a 50-year-old symptomatic male patient in Dhaka, Bangladesh (SARS-CoV-2 strain G039392) and the strain was found as 99.9% identical to the UK variant B.1.1.7 [12]. In this study, six proteins (ORF10, ORF7b, ORF7a, ORF6, membrane glycoprotein, and envelope protein) were randomly selected from the SARS-CoV-2 Bangladeshi strain G039392 regarding characterization, so that more detailed studies could elucidate their structures and provide insights into the possible functions for the selected proteins. Therefore, the major purpose of this analysis is to learn more about these proteins like the assessment of a.a composition, the energy level of chemical bonds, hydropathicity, etc. through bioinformatics approaches which could provide insight into probing novel functions regarding virulence of Covid-19. Moreover, structural prediction of 2D and 3D SARS-CoV-2 protein models could give further way to docking molecular components which can optimize devastating viral properties of this particular virus. Thereby, this study utilizes strictly bioinformatics approaches to theoretically characterize, classify, and construct the putative structure of selected 6 proteins in SARS-CoV-2 Bangladeshi strain G039392.

## 2. Materials and Method

### 2.1 Sequence Alignment and Phylogenetic Analysis

The reference sequence corresponding to the selected six proteins (ORF10, ORF7b, ORF7a, ORF6, membrane glycoprotein, and envelope protein) in SARS-CoV-2 strain G039392, along with other 9 variants from 9 different countries, were acquired from NCBI’s Protein Database. Sequences were aligned using MUSCLE on the MEGA-X-V10.1.7 software [13,14]. The neighbor-joining method was implemented by maintaining other default settings. Alignments reliability was measured by overall mean distance (≤ 0.7 is reliable) and determined using p-distance substitution model [15]. The protein trees were constructed using the neighbor-joining method and visualized on MEGA-X-V10.1.7. The phylogeny trees were tested using the bootstrap method.

### 2.2 Protein Characterization

Phosphorylation sites were detected by DEPP server of PONDR® [16]. ProtScale was used to generate hydrophobicity plot and ProtParam to determine the grand average of hydrophobicity (GRAVY) [17,18]. Also, the protein disorder predictions were performed using PONDR® (Predictor of Natural Disordered Regions) VLS2, XL1 [19]. Moreover, amino acid compositions and aliphatic index were analyzed employing ProtParam. Finally, the protein-binding propensities of the interacting residues were evaluated using SCRIBER [20].

### 2.3 Protein Secondary Structure Prediction, 3D Modeling, Evolution and Validation

NetSurfP v2.0 server was employed to predict the protein secondary structure [21]. The predictions of transmembrane helix (TH) were performed by TMHMM [22] and Phobius [23] by averaging the predictions and the most constant range of scores were utilized for analysis. The web server IntFOLD was used to make use of an *ab initio* modeling for constructing the selected proteins [24]. According to the IntFOLD’s quality and confidence scoring, the models were evaluated and utilizing the 3Drefine web-server, the best model was then refined [25]. The maximum QMEAN Z-score [26] and Ramachandran plot [27] were considered as most favorable among the five generated post-refinement models. Both UCSF Chimera and PyMol were used to visualize the most favorable 3D protein model [28,29]. The hydrophobicity surfaces were created according to the Kyte-Doolittle scale [30].

## 3. Results

### 3.1 Sequence Alignment and Phylogenetic Analysis

In phylogenetic tree, the overall mean distance of ORF10 is 0.01, which is corresponding to almost 99.9% identity for the entire alignment (Figure 1G). The ORF10 protein from the strain of Spain has shown difference at the 30^th^ position which is Leu (L) rather than Val (V) (Figure 1A). So, the height of the conserved region is from 1 to 29 residues. While, for ORF7b, ORF7a and ORF6, membrane glycoprotein, envelope protein, the mean distance is 0.00 along with 100% conserved regions which are correspondences for the entire alignment (**Figure 1**).

**Figure 1.**
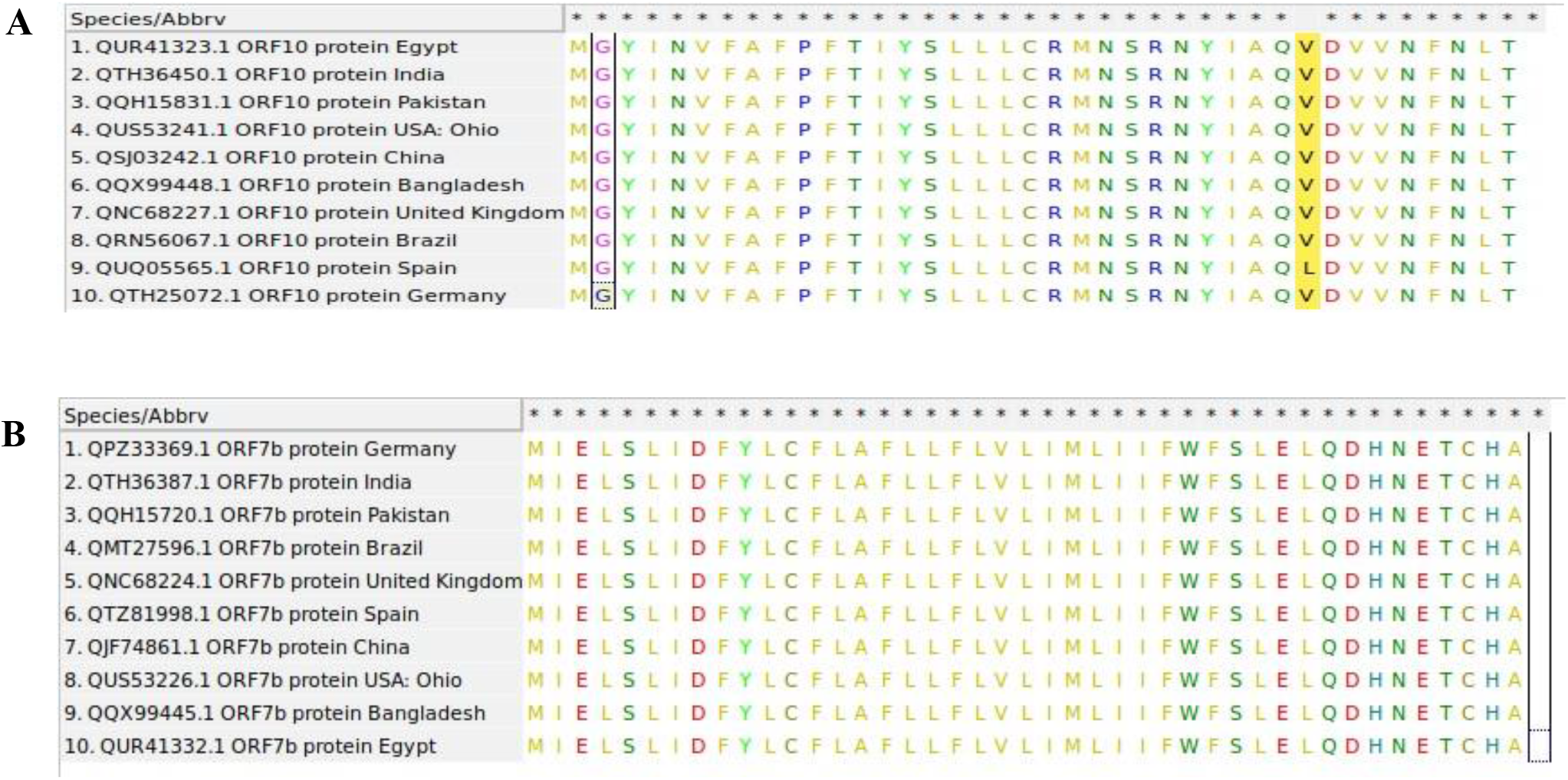
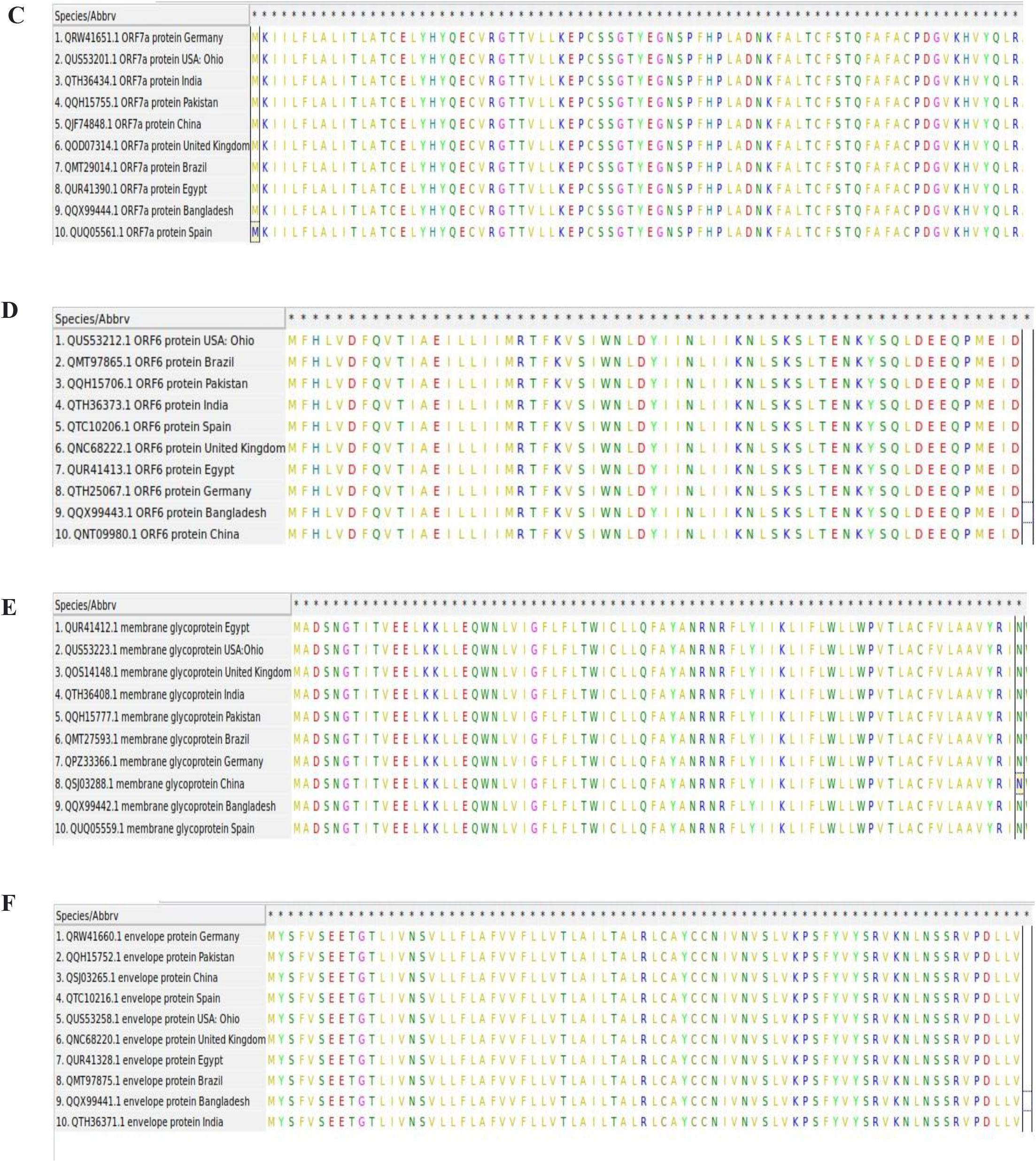
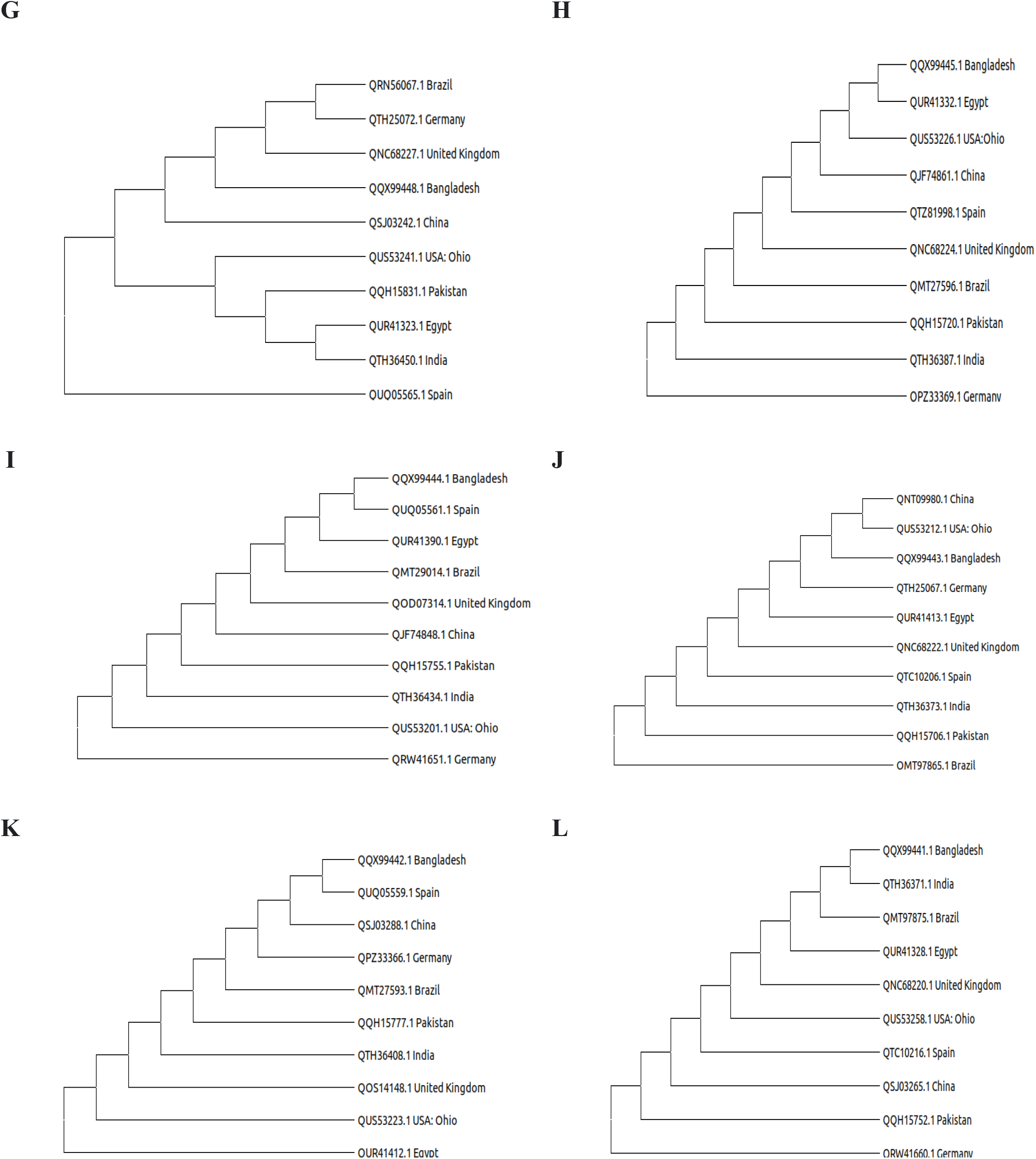
Multiple sequence alignment and Phylogenetic analysis of six different protein sequences depicting evolutionary relationships with SARS-CoV-2 varieties of 10 different Countries. **1A-F**. Sequence alignment of ORF10, ORF7b, ORF7a and ORF6, membrane glycoprotein, and envelope protein respectively. **1G-L**. Neighbor joining phylogenetic tree of ORF10, ORF7b, ORF7a and ORF6, membrane glycoprotein, and envelope protein respectively. All proteins appear closely related to the proteins of different variants of different countries.

### 3.2 Phosphorylation

Single phosphorylation site (phosphorylated serine) was identified in ORF7a consist of 14.29%, whereas other proteins did not show any Phosphorylation sites (**Table 1**).

**Table 1.**
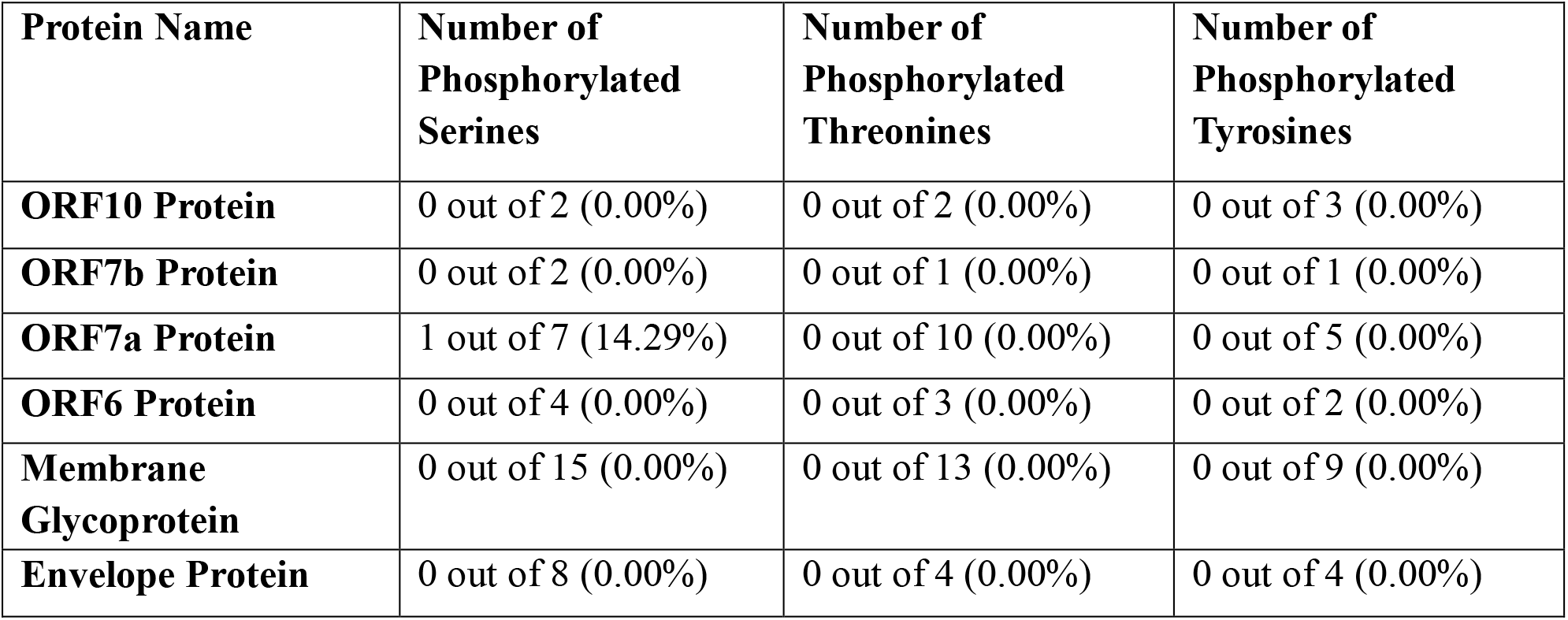
Phosphorylation sites of the selected proteins

### 3.3 Hydropathicity

In ORF10 protein, the hydrophobicity plot exposed two hydrophobic regions spanning residues 3-20 and 28-35 along with two hydrophilic regions; residues 21-27 and a residue of 36 (**Figure 2A**).There are single hydrophilic and hydrophilic regions spanning residues 3-32 and 33-41, respectively in ORF7b protein (**Figure 2B**). In case of ORF7a protein, hydrophobic regions are 3-16, 25,28-31, 47-48, 54-61, 63-67, 69, 72, 84-88, 98-115 and the hydrophilic regions are 17-24, 26-27, 32-46, 49-53, 62,68, 70-71, 76-83, 89-97, 116-119 with the neutral regions 73, 75 (**Figure. 2C**). In ORF6 protein, the hydrophobic regions are 3-7, 9-20, 23-28, 31-39, 42, hydrophilic regions are 8, 21-22, 29, 40-41, 43-59 with a neutral position 30 (**Figure 2D**). Moreover, for membrane glycoprotein, 8-11, 15, 21-39, 46-71, 74-75, 77-102, 104, 110, 117-122, 124, 126-132, 134, 137-146, 149-151, 168-171, 182-183, 189, 193-195, 217-220 are hydrophobic positions, while 3-7, 12-14, 16-20, 40-45, 72-73, 76, 103, 105-109, 111-116, 123, 125, 133, 135-136, 147-148, 152-167, 173-180, 184-188, 190-192, 196-216 are hydrophilic position and neutral places are 172, 181(**Figure 2E**). Whereas, in case of envelop protein, hydrophobic region residues are 3-5, 11-54, 56-58,60, 72-73 and hydrophilic region residues are 6-10, 55, 59, 61-71(**Figure 2F**). The GRAVY scores are 0.64, 1.45, 0.32, 0.23, 0.45 and 1.13, respectively for ORF10, ORF7b, ORF7a, ORF6, membrane glycoprotein and envelope protein (**Figure 2G**).

**Figure 2.**
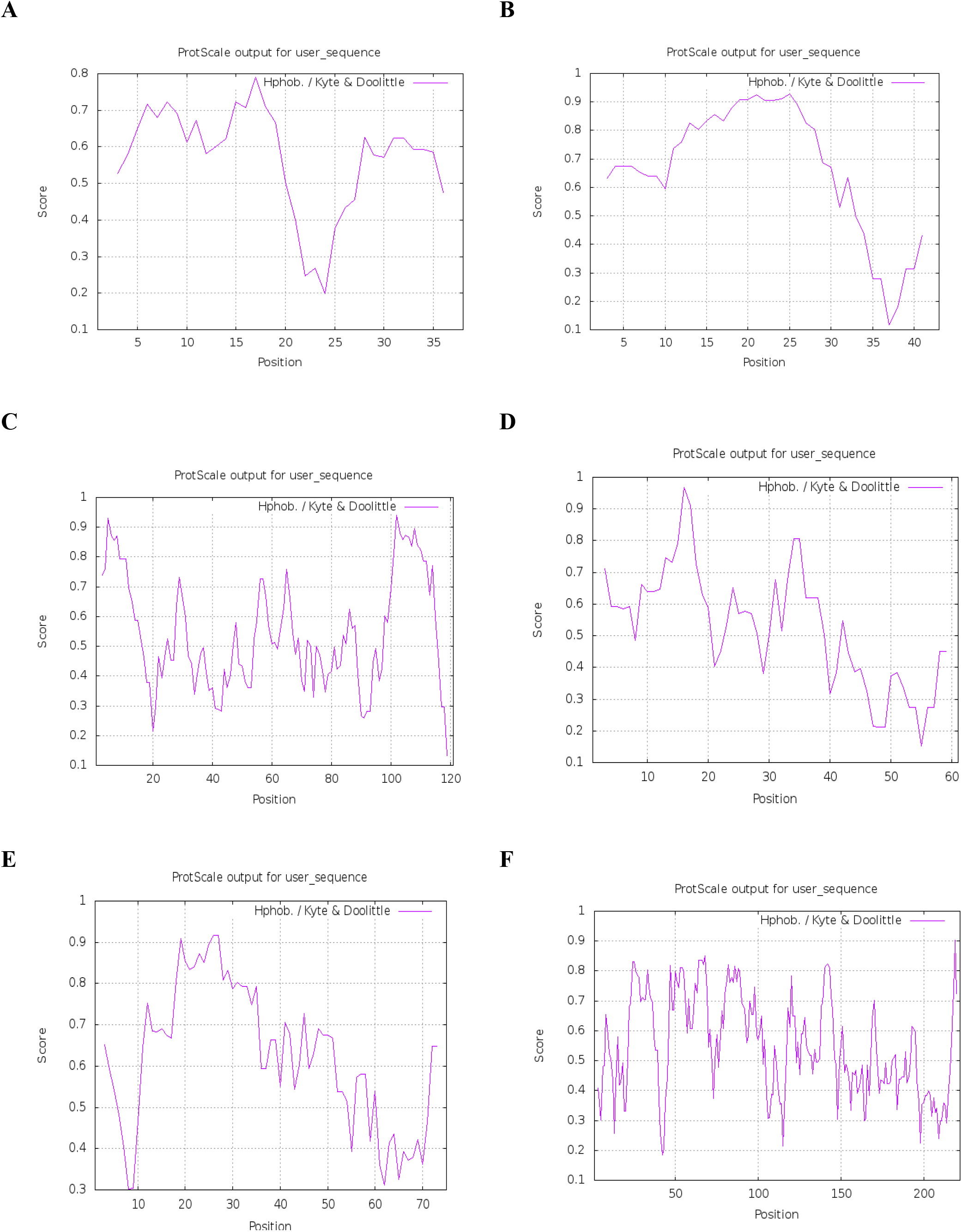
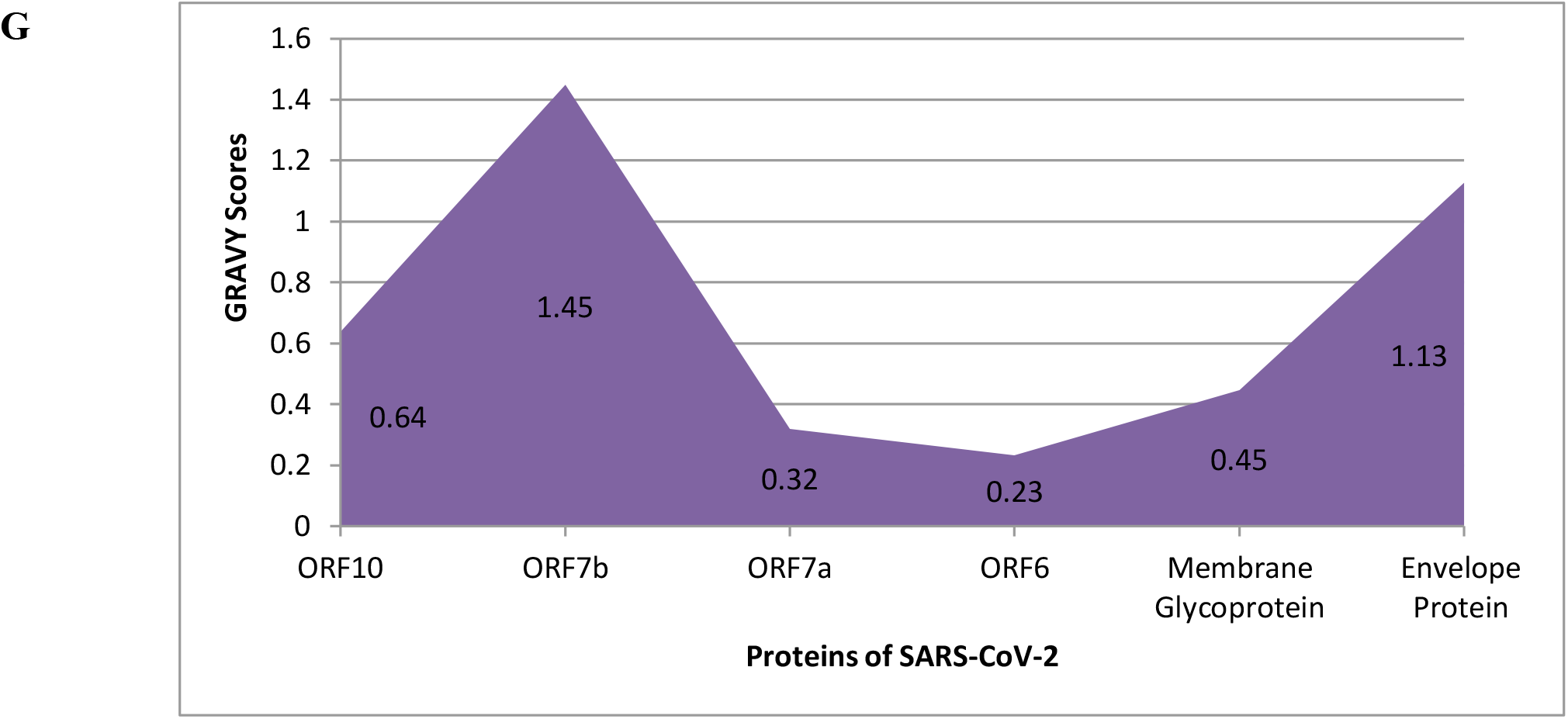
Hydrophobicity plot and GRAVY Scores of selected 6 proteins. **2A-F**. Hydrophobicity plot of ORF10, ORF7b, ORF7a and ORF6, membrane glycoprotein, and envelope protein respectively. The hydrophobicity plots were generated according to the Kyte-Doolittle hydropathy plots. **2G**. GRAVY Scores of ORF10, ORF7b, ORF7a, ORF6, membrane glycoprotein, and envelope protein. The numerical values for each score displayed above are their corresponding box. The proteins are recognized as mostly hydrophobic.

### 3.4 Protein Disorder

The protein disorder plot indicated that the disorder scores were higher for C-terminal half than N-terminal half for almost all selected proteins (**Figure 3**). Almost all proteins showed protein disorder scores indicating moderate flexible to highly flexible residues. However, membrane glycoprotein is more disordered compared to other proteins. Also, no protein revealed scores of ≤ 0.1 indicating rigidity.

**Figure 3.**
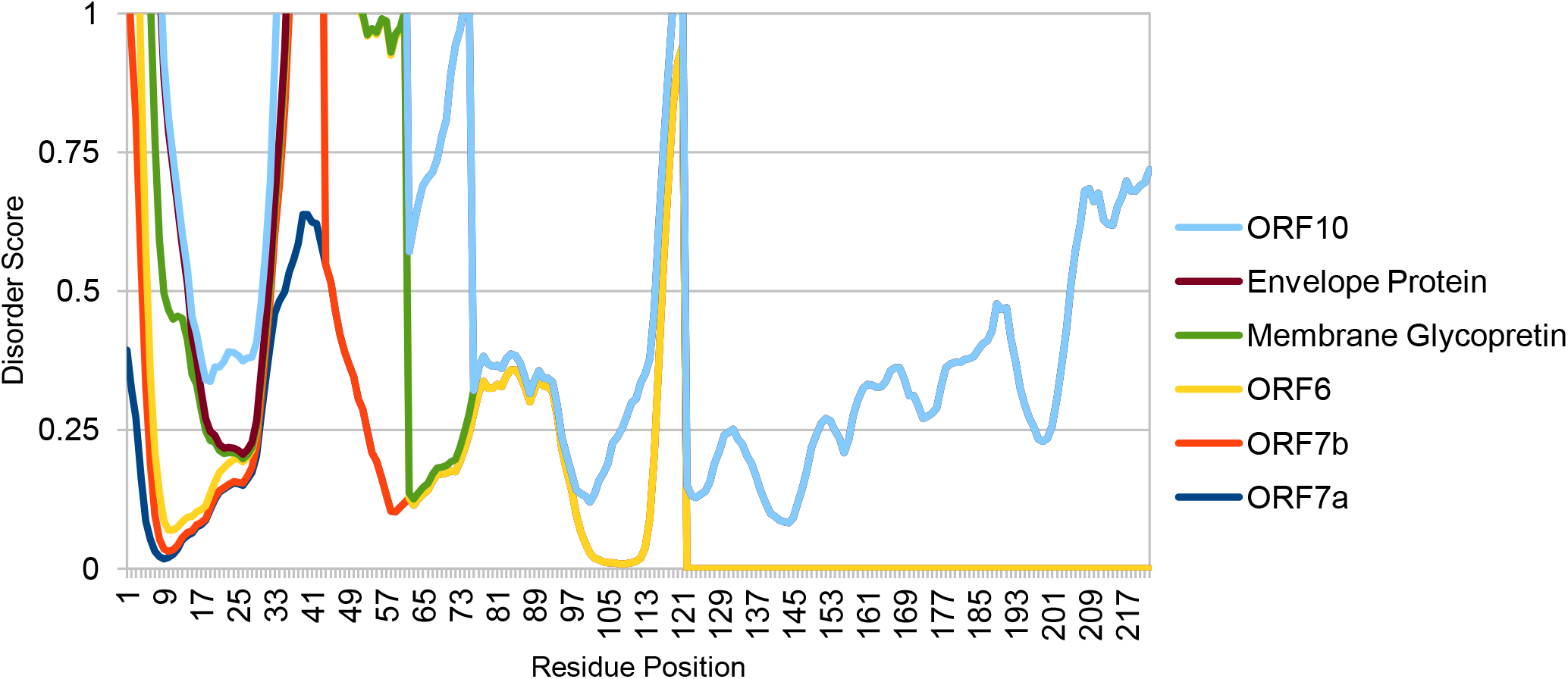
Per-residue disorder plot for ORF10, ORF7b, ORF7a, ORF6, membrane glycoprotein, and envelope protein of SARS-CoV2. All proteins found as highly flexible and ordered. Scores ≥ 0.5 indicate disorder residues, while scores within 0.25-0.5 and 0.1-0.25 suggest highly flexible and moderate flexible residues. Scores ≤ 0.1 indicate rigidity.

### 3.5 Amino Acids Composition and Protein-Binding Propensity

The ORF10 protein consists of the highest percentage of asparagines (N), where, in case of ORF7b, ORF7a, membrane glycoprotein, and envelope protein, leucine (L) is presented in the maximum percentage. Moreover, for ORF6, it was isoleucine (Ile). The overall amino acid composition of all 6 proteins has been represented in **Table 2**. The binding propensity is important to influence electrostatic and aromatic interactions and also it is extremely varied with the amino acid residues. Several fluctuations have been observed in protein-binding propensity in both C-terminal half residues and N-terminal half residues (**Supplementary Figure 1**).

**Table 2.** Amino Acid composition of ORF10, ORF7b, ORF7a, ORF6, membrane glycoprotein, and envelope protein in percentage (%)

| Amino Acid | ORF10 | ORF7b | ORF7a | ORF6 | Membrane Glycoprotein | Envelop Protein |
| --- | --- | --- | --- | --- | --- | --- |
| Ala (A) | 5.3% | 4.7% | 7.4% | 1.6% | 8.6% | 5.3% |
| Arg (R) | 5.3% | 0.0% | 4.1% | 1.6% | 6.3% | 4.0% |
| Asn (N) | 13.2% | 2.3% | 1.7% | 6.6% | 5.0% | 6.7% |
| Asp (D) | 2.6% | 4.7% | 1.7% | 6.6% | 2.7% | 1.3% |
| Cys (C) | 2.6% | 4.7% | 5.0% | 0.0% | 1.8% | 4.0% |
| Gln (Q) | 2.6% | 2.3% | 4.1% | 4.9% | 1.8% | 0.0% |
| Glu (E) | 0.0% | 7.0% | 6.6% | 8.2% | 3.2% | 2.7% |
| Gly (G) | 2.6% | 0.0% | 3.3% | 0.0% | 6.3% | 1.3% |
| His (H) | 0.0% | 4.7% | 2.5% | 1.6% | 2.3% | 0.0% |
| Ile (I) | 7.9% | 11.6% | 6.6% | 16.4% | 9.0% | 4.0% |
| Leu (L) | 10.5% | 25.6% | 12.4% | 13.1% | 15.8% | 18.7% |
| Lys (K) | 0.0% | 0.0% | 5.8% | 6.6% | 3.2% | 2.7% |
| Met (M) | 5.3% | 4.7% | 0.8% | 4.9% | 1.8% | 1.3% |
| Phe (F) | 10.5% | 14.0% | 8.3% | 4.9% | 5.0% | 6.7% |
| Pro (P) | 2.6% | 0.0% | 5.0% | 1.6% | 2.3% | 2.7% |
| Ser (S) | 5.3% | 4.7% | 5.8% | 6.6% | 6.8% | 10.7% |
| Thr (T) | 5.3% | 2.3% | 8.3% | 4.9% | 5.9% | 5.3% |
| Trp (W) | 0.0% | 2.3% | 0.0% | 1.6% | 3.2% | 0.0% |
| Tyr (Y) | 7.9% | 2.3% | 4.1% | 3.3% | 4.1% | 5.3% |
| Val (V) | 10.5% | 2.3% | 6.6% | 4.9% | 5.4% | 17.3% |
| Pyl (O) | 0.0% | 0.0% | 0.0% | 0.0% | 0.0% | 0.0% |
| Sec (U) | 0.0% | 0.0% | 0.0% | 0.0% | 0.0% | 0.0% |

### 3.6 Aliphatic Index and Transmembrane Helix

Aliphatic index values of more than 100 indicated that these proteins are highly thermo-stable over a wide temperature assortment. For ORF10, ORF7b, ORF7a, ORF6, membrane glycoprotein, and envelope protein, the aliphatic index values found were 107.63, 156.51, 100.74, 130.98, 120.86, and144 respectively (**Figure 4A**). Transmembrane helices of less than one indicated these helices are less likely to interact with membrane lipids. For ORF10 and ORF7a, TH predicted spanning residues are 3-29 and 6-33, respectively. In case of ORF7b, TH predicted residues are 4-23 and 93-119. Furthermore, for ORF6 and envelope protein, the predicted TH spanning residues are 5-38 and 11-61. Finally, in case of membrane glycoprotein, the transmembrane helix residues are 18-59 and 61-105. The representation is in **Figure 4B**.

**Figure 4.**
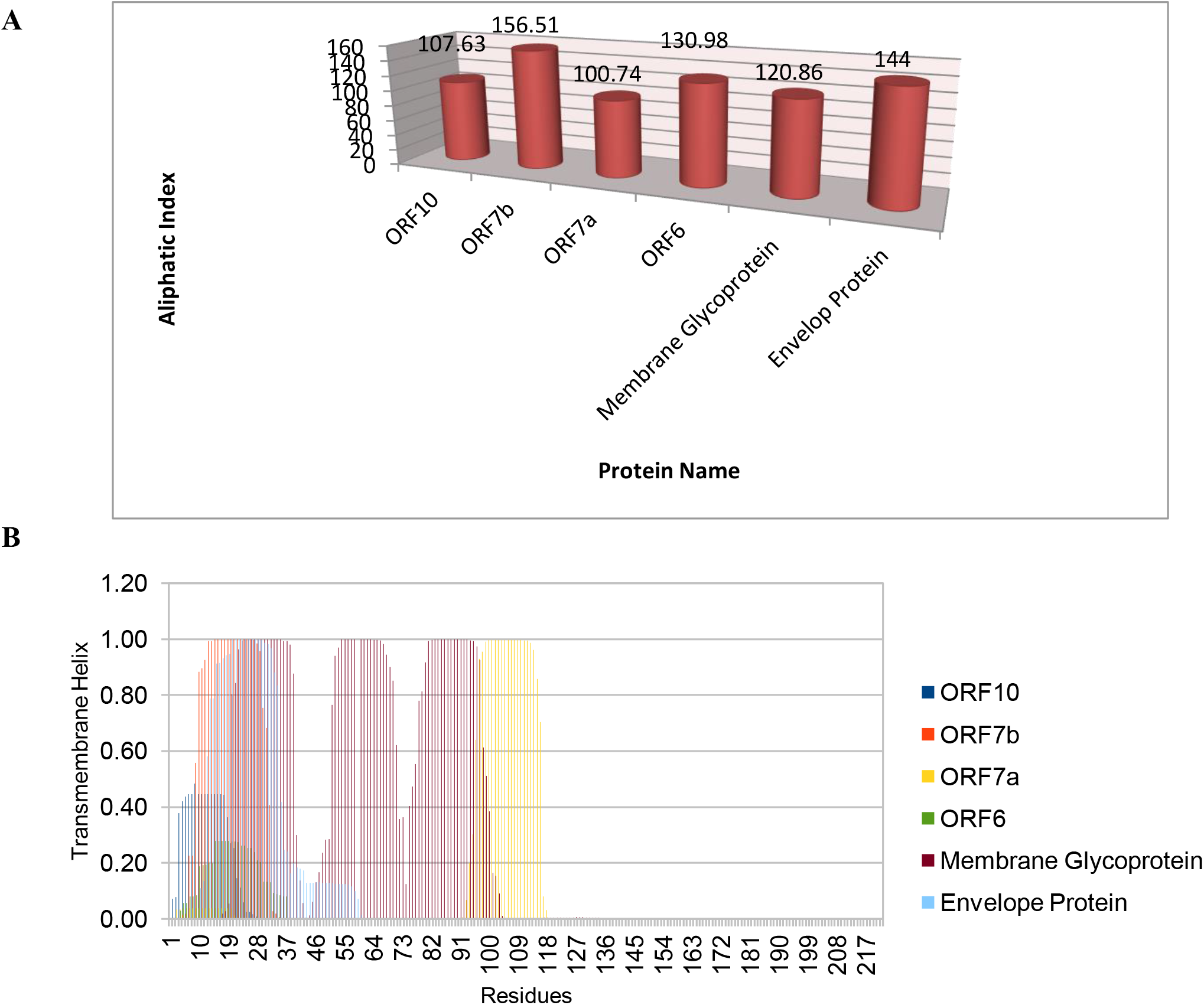
Aliphatic indexes and transmembrane helix prediction scores of ORF10, ORF7b, ORF7a, ORF6, membrane glycoprotein, and envelope protein respectively (**4A-B**). All proteins showed aliphatic index values of more than 100 indicated that they are highly thermo-stable and transmembrane helices of less than 1.

### 3.7 Protein Secondary Structures

In respect to ORF10, ORF7b and envelope protein, α-helix spanning residues are 11-21, 4-35 and 4-64, respectively (**Figure 5A, 5B, 5F**). Additionally, in ORF6, the α-helix spanning residues are 4-21, 26-27, 29-44, 48-51 (**Figure 5D**). In case of ORF7a, the α-helix and β–sheet spanning residues are 90-96, 99-100 and 28-33, 40-41, 53-66, 72-79, respectively (**Figure 5C**). The membrane glycoprotein has the α-helix and β–sheet spanning residues of 10-19, 22-36, 40-70, 75-106, 161-163 and 112, 118-123, 128-132, 139-146, 148-151, 154-159, 167-172, 175-185, 193-201, respectively (**Figure 5E**).

**Figure 5.**
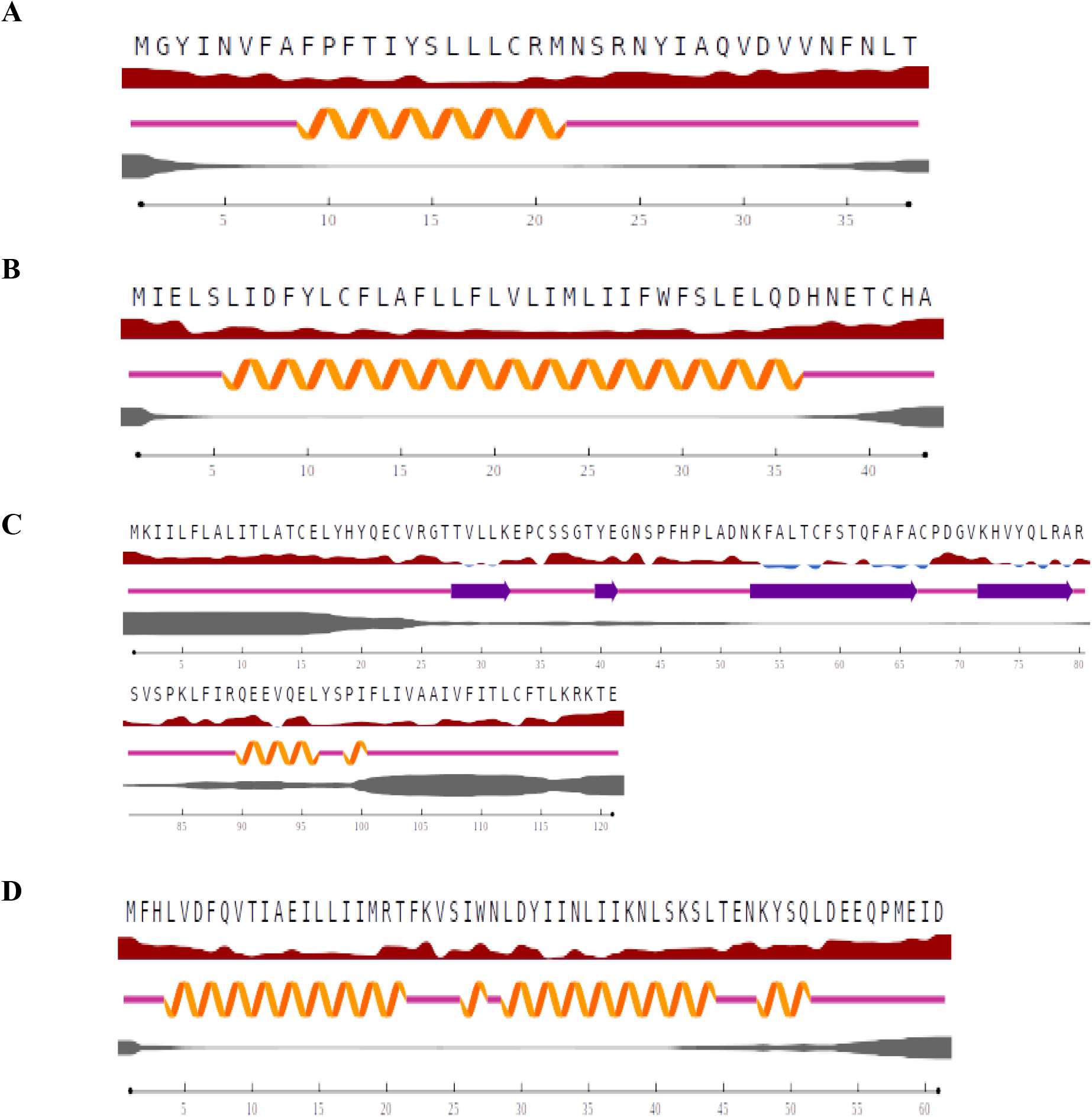
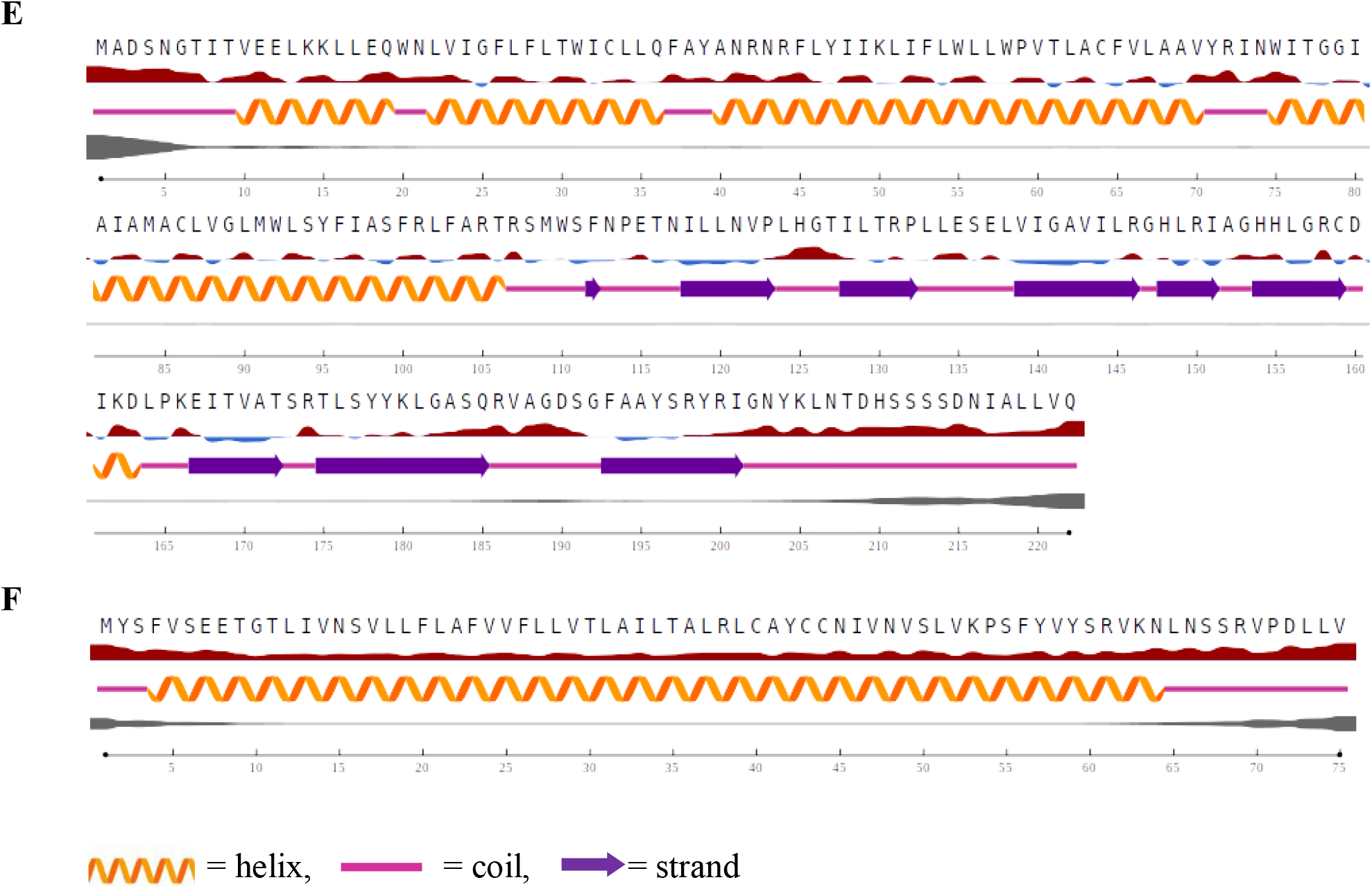
Secondary structure prediction of 6 selected proteins. **5A-F**. Secondary structure of ORF10, ORF7b, ORF7a, ORF6, membrane glycoprotein, and envelope protein respectively.

### 3.8 Protein Modeling and Validation

Initially, the models having low P-values and high-quality scores were subjected to refinement which was yielded by the IntFOLD web-based server. Then the selection was done according to the QMEAN Z score and Ramachandran plot score (**Table S1** and **Figure 6A-F**). QMEAN Z score and Ramachandran plot score table were added as supplementary table 1 (**Table S1)**. In response to hydrophobic and hydrophilic properties, the majority of the proteins surfaces were found as hydrophobic (**Figure 6G-L**). The Ramachandran plot score details for all the proteins were found more than 90% except for membrane glycoprotein which is 87.6% (**Figure 6M-R**).

**Figure 6.**
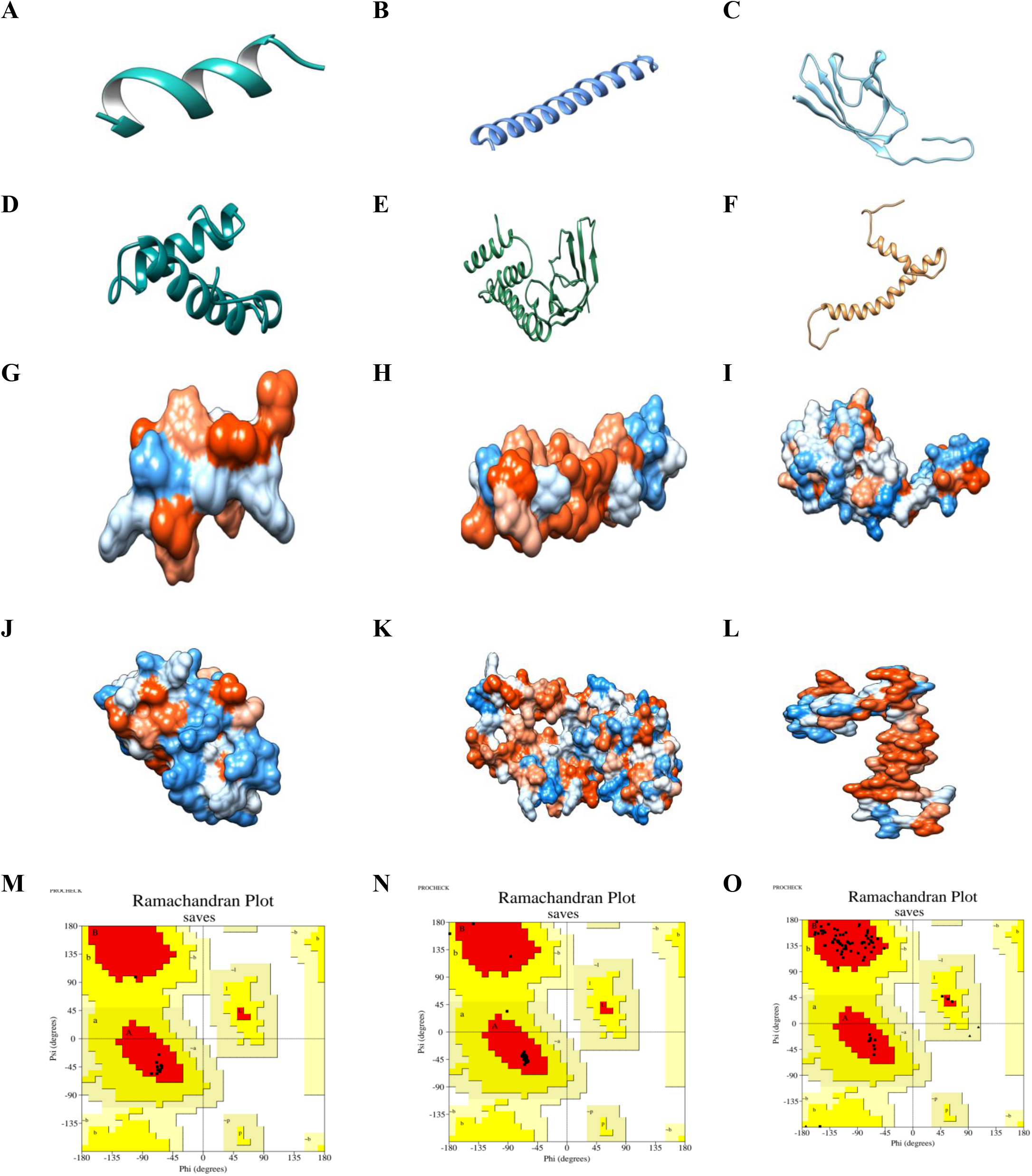
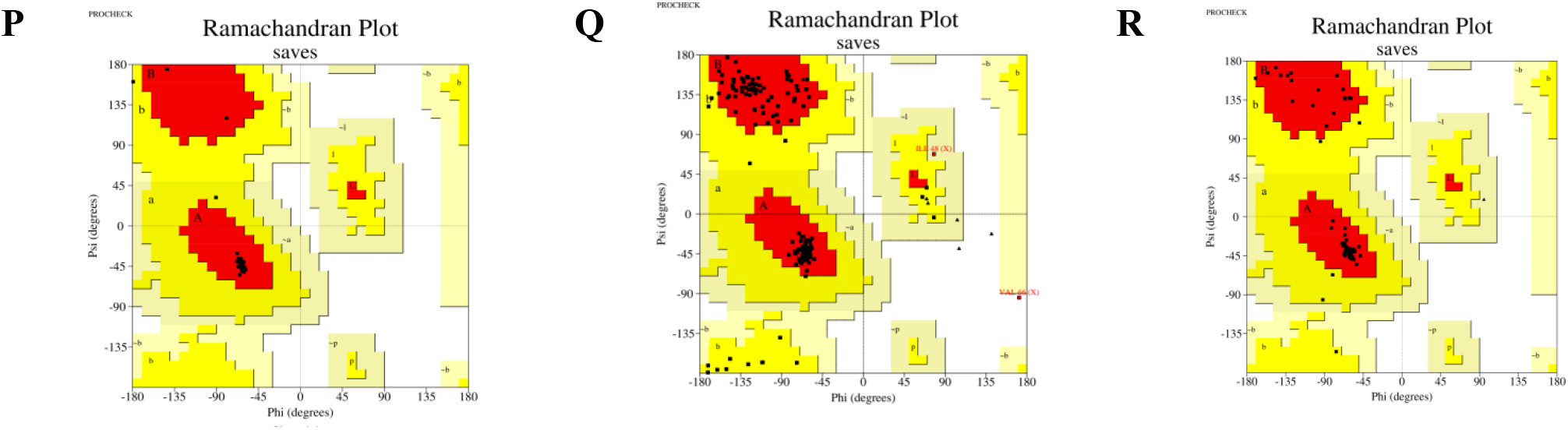
Protein modeling and hydrophobicity surface 3D map of the selected 6 proteins. **6A-F**. Ribbon diagram of ORF10, ORF7b, ORF7a, ORF6, membrane glycoprotein, and envelope protein respectively. **6G-L**. Hydrophobicity surface map of ORF10, ORF7b, ORF7a, ORF6, membrane glycoprotein, and envelope protein respectively. All 6 protein models are found as highly flexible and stable. The blue color represents hydrophilic regions and the orange color expresses hydrophobic regions. Where, the whitish-blue color indicates semi-hydrophobic/hydrophilic character. **6M-R**. Representation of Ramachandran plots of ORF10, ORF7b, ORF7a, ORF6, membrane glycoprotein, and envelope protein, respectively.

## 4. Discussion

At most, the phylogenetic data in this study proposes the ORF10 protein in Spain is most distantly related to all other ORF10 proteins. Whereas, the high similarity was detected in the other remaining selected ORF10 proteins. While, for ORF7b, ORF7a and ORF6, membrane glycoprotein, envelope protein, there was no distant relationship among the stains from the selected different countries. All these proteins showed 100% conserved region, thus, mutations in these regions were not created along with revealed that these proteins shared a strong phylogenic relationship with their common ancestors in the past. Conserved regions of ORF7b, ORF7a and ORF6, membrane glycoprotein, and envelope protein could be playing a fundamental role in the assembly of particular proteins, formation of protein structure and/or generating virulent functions by facilitating precise protein interactions. Although, defining relationships between specific sequences is not entirely possible when based solely on sequence data [31]. We can predict that the proteins here are highly ordered as intrinsically disordered proteins have a tendency to be phosphorylated that leads to disorder-to-order along with order-to-disorder transitions [32].

Phosphorylation controls the function of the protein and cell signaling by changing conformational shape in the phosphorylated protein which maintains the catalytic property of the particular protein. Thus, activation or inactivation of proteins depends on phosphorylation [33]. Phosphorylation site prediction of selected 6 proteins conceded that phosphorylated serine, threonine or tyrosine was mostly not present though ORF7a was showed a single phosphorylated serine. In every conceivable way, the phosphorylation of a distinct protein is able to modify its activities which include inflection of protein’s intrinsic biological property, proper sub-cellular location, docking with other related proteins and half-life. It also decides the level and period of a response given by a protein which acts as input to signal integration [34]. Moreover, sites of phosphorylation are more prone to be evolutionary conserved than other interfacial residues [35].

The purpose of the hydropathy index of amino acids is mainly to predict the function of a structurally or functionally unknown protein. The distribution of hydropathy clusters in a particular protein appears to recommend that these cluster location is principally conserved in a given group of proteins [36]. In this present study, selected 6 proteins showed hydropathy index which tended to be more hydrophobic. The literature revealed that hydrophobic proteins are more soluble and for this reason, they can function in an independent manner by avoiding undesirable interactions with watery molecules. In addition to that, these proteins are vital for protein folding which keeps it more stable and biologically active [37].

Protein disorder predictions revealed that almost all proteins showed protein disorder scores indicating moderate flexible to highly flexible residues and no protein revealed scores which indicates rigidity. This result coincides with the result presented by [18]. However, membrane glycoprotein was more disordered compared to other proteins in this study. Protein disorder predictions are an enormous challenge in structural <u>proteomics</u> and subsequently its function prediction including identification of those proteins that are unstructured either partially or wholly. Disordered regions present in specific proteins could contain short linear peptide motifs which may later play a significant role in protein function. After predicting, avoidance of prospective disordered regions in protein can augment expression, proper foldability, and stability of that expressed protein [38].

Protein binding propensity augments the knowledge of protein-protein interactions, docking, and annotation of functional properties of that protein at the molecular level [20]. Also, a high aliphatic index resembles to rise of the thermostability of globuler proteins [39]. All 6 selected proteins showed aliphatic index of more than 100 which indicates these proteins are highly thermo-stable over a wide temperature assortment. In addition, all 6 selected proteins showed transmembrane helixes which are less than 1 and transmembrane helixes have immense importance in the study of membrane proteins [40].

Due to the significance of structural class prediction of protein, diverse major efforts have been employed to discover a prediction model that establishes the structural class and predicts protein secondary structure depending on the sequences of protein [41,42]. The prediction of secondary structures for six selected proteins revealed that each ORF7a protein and membrane glycoprotein has one α-helix and one β-strand. The structural class is one of the most imperative features for its vital task in the analysis of protein function, prediction of the rate of protein folding nature, and, also, execution of a suitable approach to uncover protein tertiary structure [43]. In this present study, the stable tertiary structure of proteins was predicted which gives the primary notion about the interaction of this protein 3D structures with enzymes or host receptors. Also, in this study, hydrophobicity surface map of particular proteins was created to distinctly show the hydrophobic or hydrophilic regions of protein.

Studying these diverse proteins of the SARS-CoV-2 virus has already yielded some clues about how they connect with the human cells but much remains to be assessed. The present study explored theoretical modeling, sequence and structure-based functional characterization of six accessory proteins. Phylogenetic analysis of these proteins exposed a close evolutionary relationship with the proteins of distant origins. Though further comprehensive assessment with broad-scale data are required to elucidate these upshots generated in this current study.

## 5. Conclusion

Communally, the present study provides an interesting basis for characterizing proteins of novel viruses theoretically and structurally. The selected proteins appear stable, ordered, hydrophobic, and also share strong phylogenetic relationships with proteins of other closely related SARS-CoV-2. Finally, the tertiary models of protein constructed in this study have higher quality and stability. This analysis can offer a foundation to perform the further analysis necessary to evaluate the biological function, interaction, and relevance to viral property of the 6 proteins in SARS-CoV-2. Perhaps, the theoretical structures would be functional for investigation of each protein interaction and their functionalities by advanced computational analysis, understanding of viral pathogenesis or to study potential vaccines and especially, to avert epidemics and pandemics.

## Data availability

The reference FASTA sequence corresponding to the selected six proteins (ORF10, ORF7b, ORF7a, ORF6, membrane glycoprotein, and envelope protein) SARS-CoV-2 strain G039392 along with other 9 variants of SARS-CoV-2 from 9 different countries were acquired and still available in NCBI’s Protein Database (https://www.ncbi.nlm.nih.gov/protein/?term=).

## Conflicts of Interest

All authors declare no conflict of interest.

## Funding

This research received no specific grant from any funding agency in the public, commercial or not-for-profit sectors.

## Supplementary material

A supplementary file is provided along with the main manuscript containing the figures of protein-binding propensity of the selected 6 proteins **(Supplementary Figure 1)** and also, a table comprising selected QMEAN Z-Scores and Ramachandran Plot Scores for protein modeling **(Table S1)**.

## Notes

https://www.ncbi.nlm.nih.gov/protein/?term

### Competing Interest Statement

The authors have declared no competing interest.

