## Supplemental Figures and Table for "Characterization and Structural Prediction of ORF10, ORF7b, ORF7a, ORF6, Membrane Glycoprotein, and Envelope Protein in SARS-CoV-2 Bangladeshi Variant through Bioinformatics Approach": Supplementry Figures and Table.pdf

**Supplementary information**

**A.**

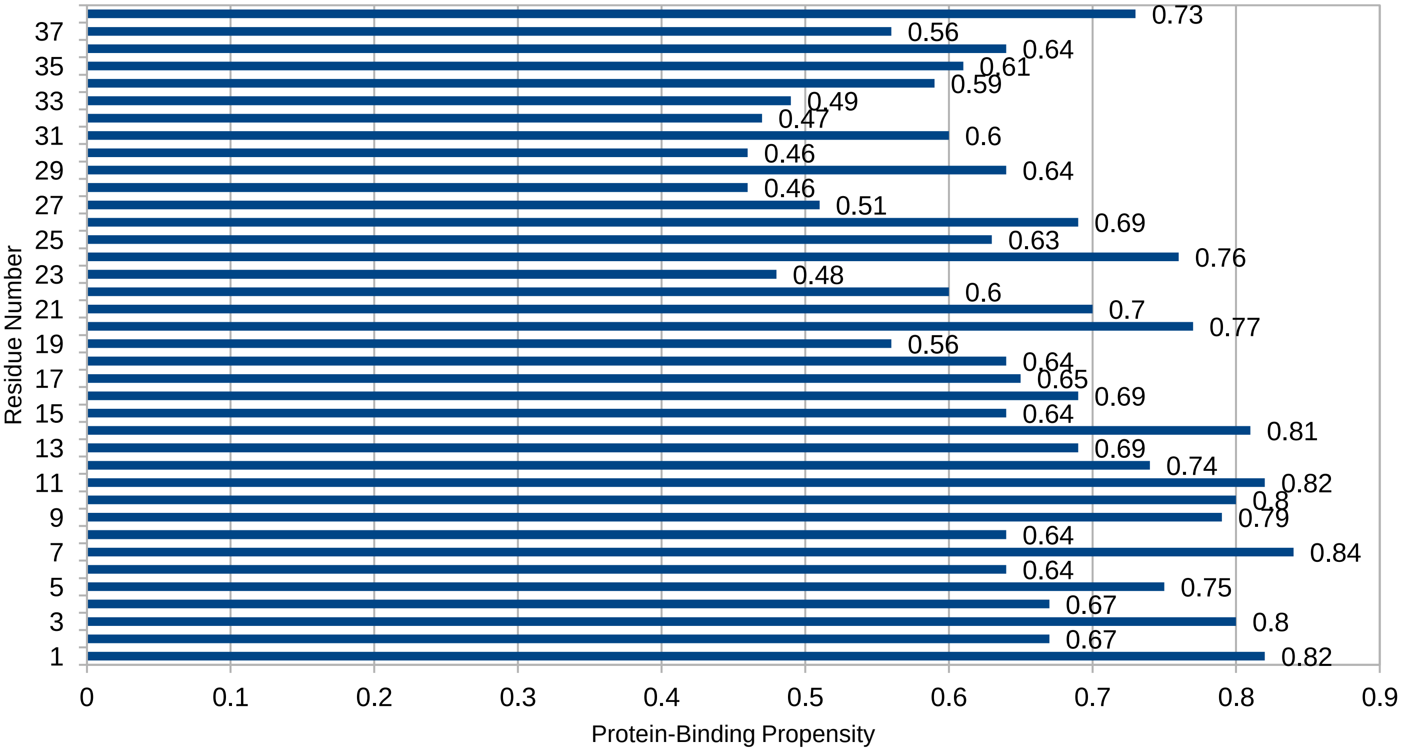


**B.**

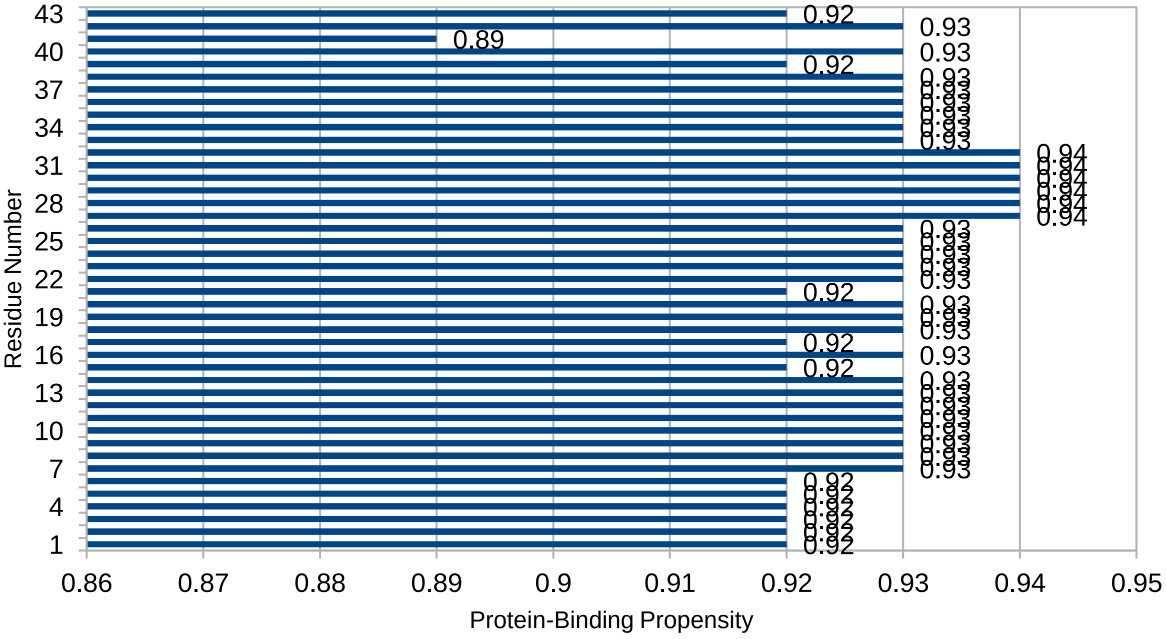

C

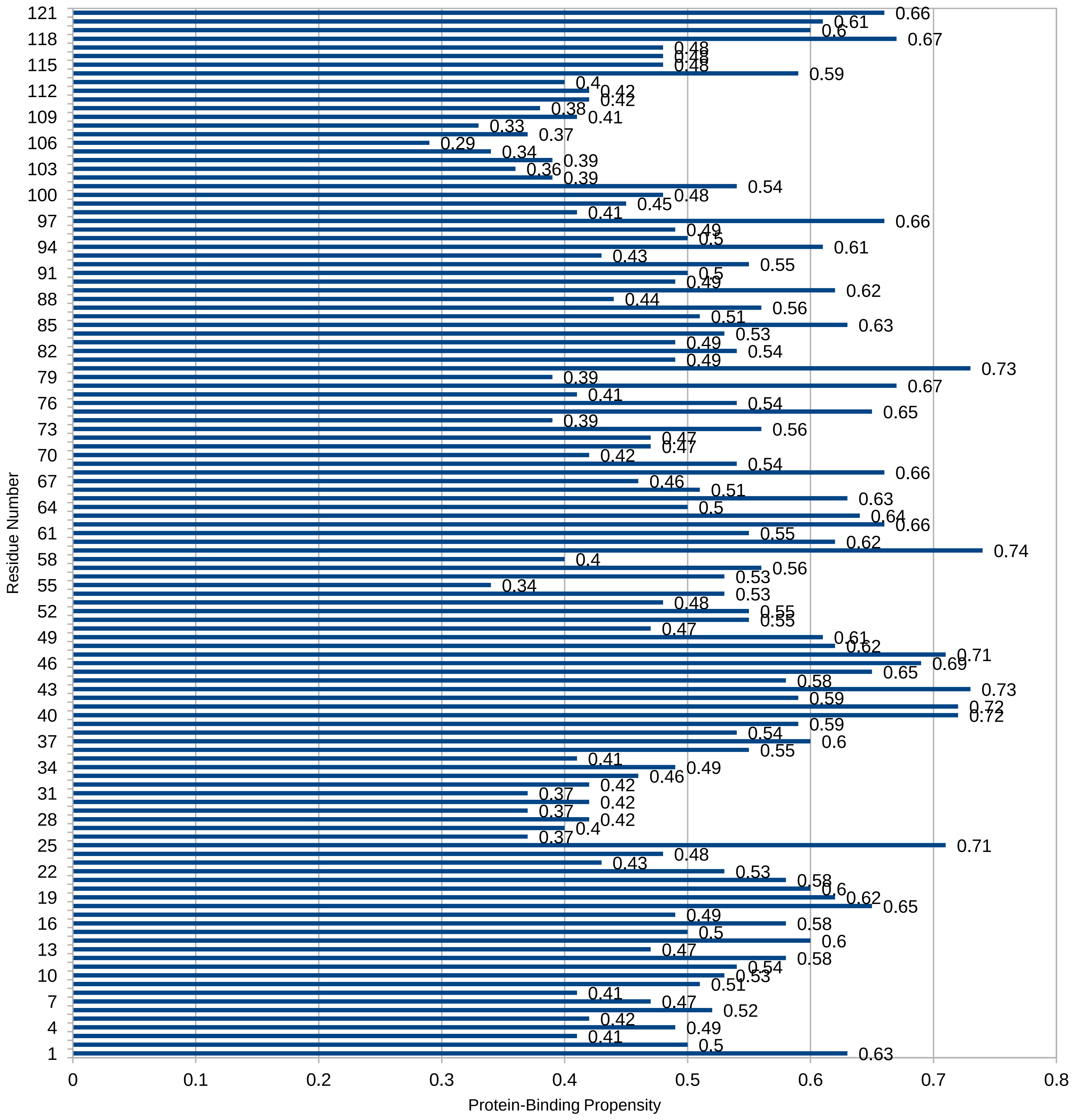

**D.**

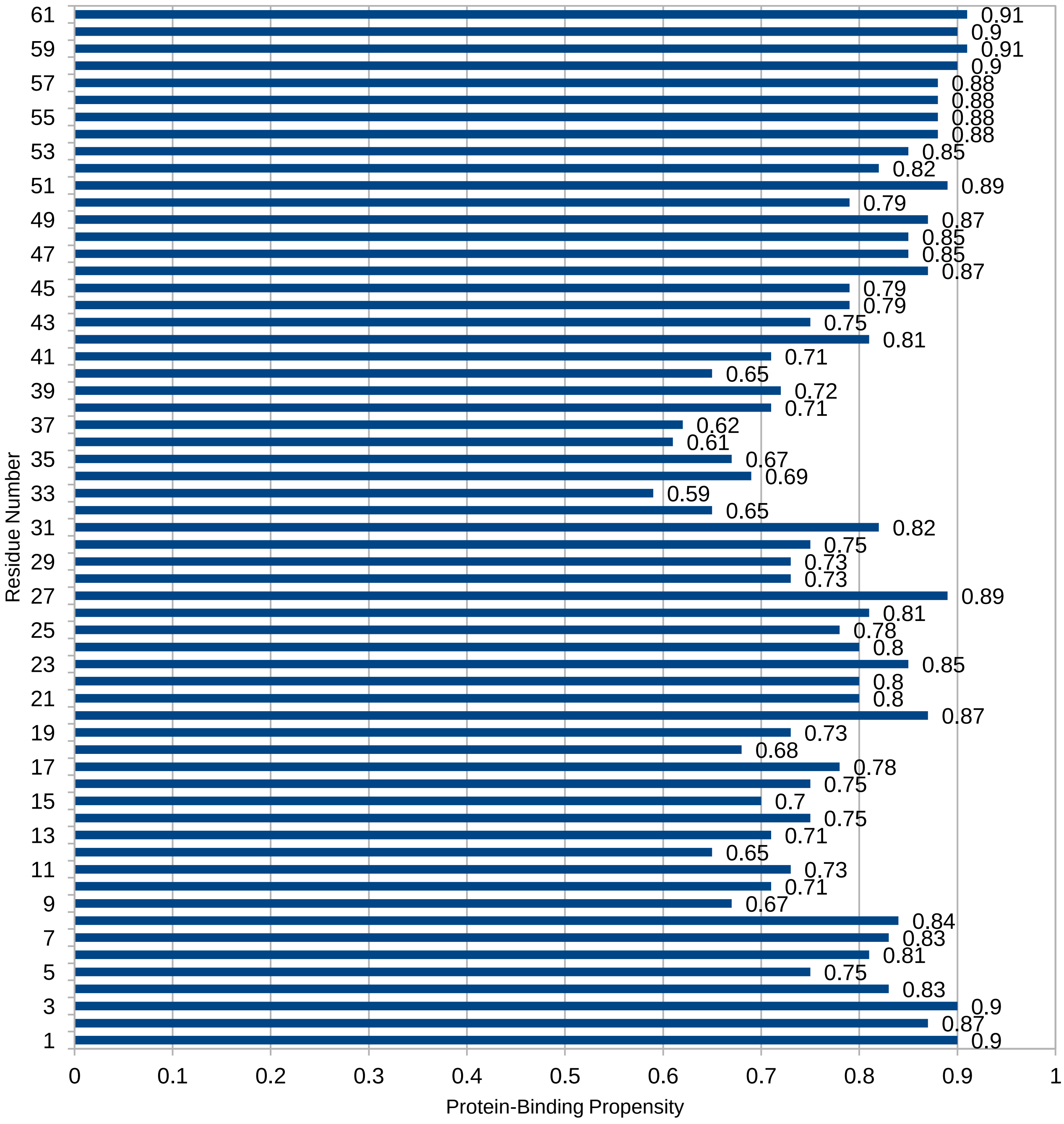

Residue Number
Protein-Binding Propensity
0
0.1
0.2
0.3
0.4
0.5
0.6
0.7
0.8
0.9
1

**E.**

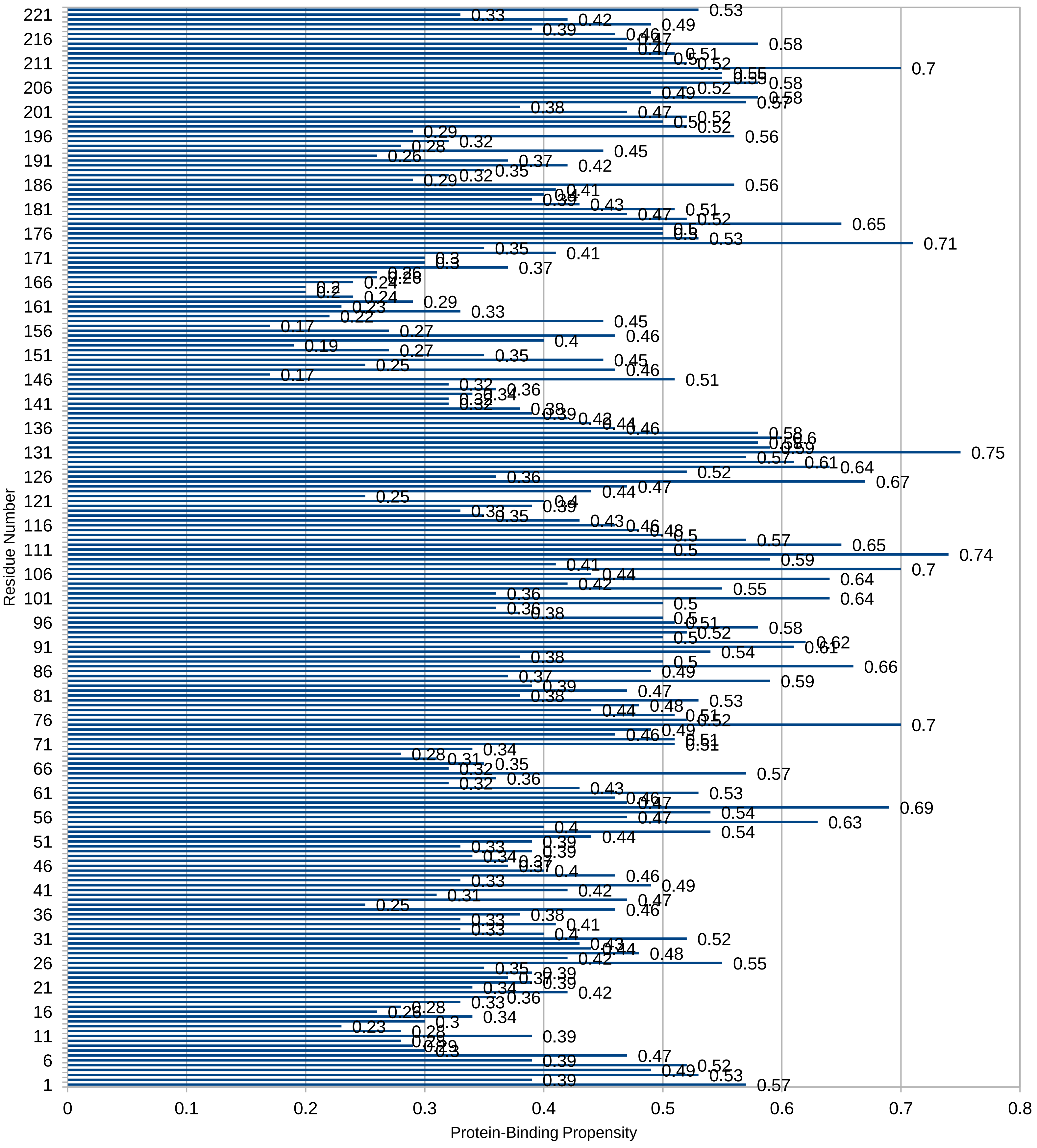

Residue Number
Protein-Binding Propensity
0
0.1
0.2
0.3
0.4
0.5
0.6
0.7
0.8
1
6
11
16
21
26
31
36
41
46
51
56
61
66
71
76
81
86
91
96
101
106
111
116
121
126
131
136
141
146
151
156
161
166
171
176
181
186
191
196
201
206
211
216
221

**F.**

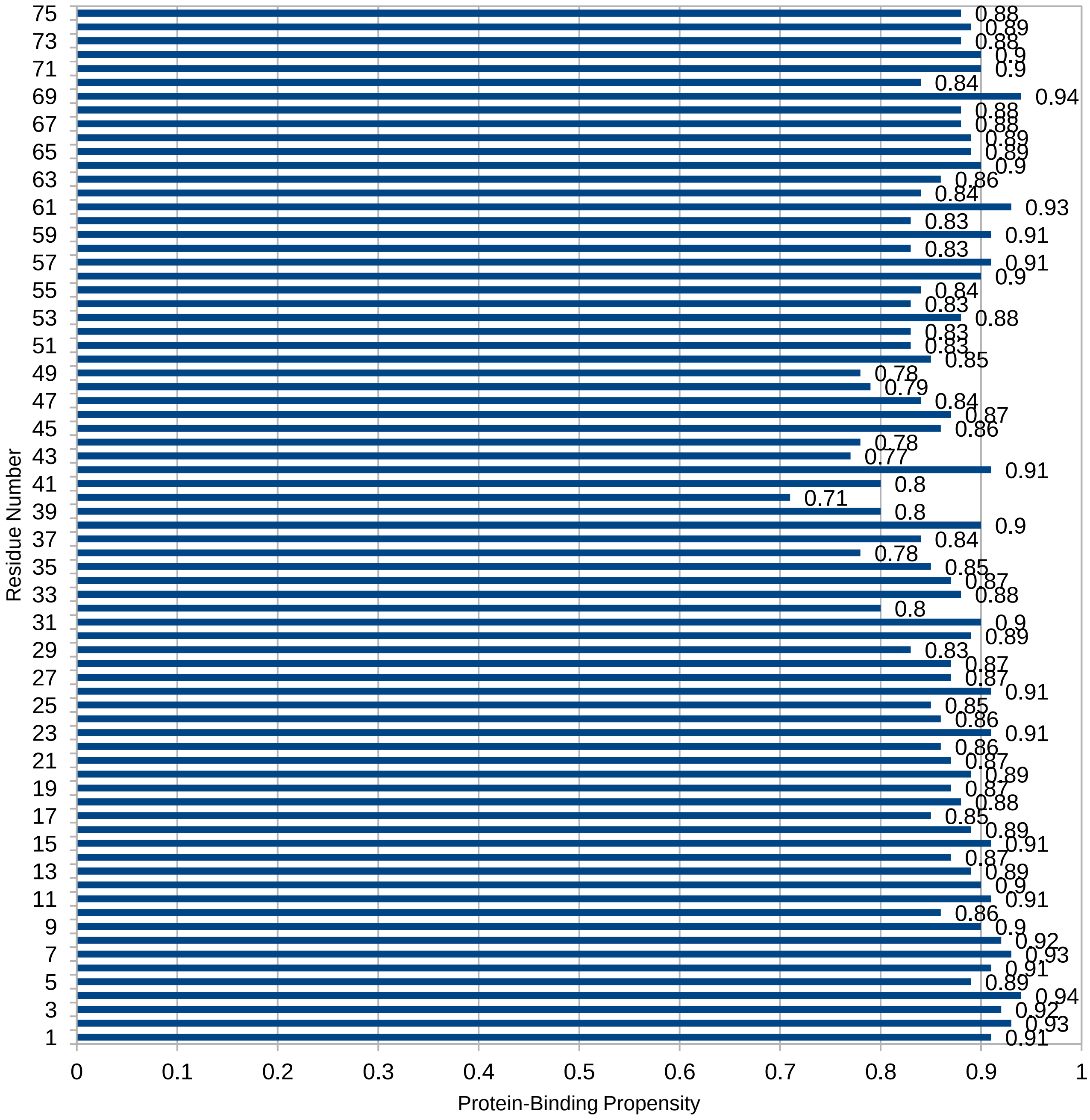


**Supplementary Figure 1.** Protein-binding propensity of the selected proteins. **A-F.** Protein-binding propensity of ORF10, ORF7b, ORF7a, ORF6, membrane glycoprotein and envelope protein, respectively.

**Supplementary Table 1**

**Table S1.** Selected QMEAN Z-Scores and Ramachandran Plot Scores for protein modeling.

| Protein Names | QMEAN Z-Scores | Ramachandran Plot Scores |
|---|---|---|
| ORF10 | -1.32 | 90.9 |
| ORF7b | -0.77 | 95.1 |
| ORF7a | -1.62 | 97.2 |
| ORF6 | -3.45 | 95.1 |
| Membrane Glycoprotein | -5.61 | 87.6 |
| Envelope Protein | -2.96 | 91.4 |
